# BGDMdocker: a Docker workflow for analysis and visualization pan-genome and biosynthetic gene clusters of bacterial

**DOI:** 10.1101/098392

**Authors:** Gong Cheng, Quan Lu, Zongshan Zhou, Ling Ma, Guocai Zhang, WU Yilei, Chao Chen

## Abstract

**Motivation:** At present Docker technology has received increasing level of attention throughout the bioinformatics community. However, its implementation details have not yet been mastered by most biologists and applied widely in biological researches. In order to popularizing this technology in the bioinformatics and sufficiently use plenty of public resources of bioinformatics tools (Dockerfile and image of scommunity, officially and privately) in Docker Hub Registry and other Docker sources based on Docker, we introduced full and accurate instance of a bioinformatics workflow based on Docker to analyse and visualize pan-genome and biosynthetic gene clusters of a bacteria in this article, provided the solutions for mining bioinformatics big data from various public biology databases. You could be guided step-by-step through the workflow process from docker file to build up your own images and run an container fast creating an workflow.

**Results:** We presented a BGDMdocker (bacterial genome data mining docker-based) workflow based on docker. The workflow consists of three integrated toolkits, Prokka v1.11, panX, and antiSMASH3.0. The dependencies were all written in Dockerfile, to build docker image and run container for analysing pan-genome of total 44 *Bacillus amyloliquefaciens* strains, which were retrieved from public? database. The pan-genome totally includes 172,432 gene, 2,306 Core gene cluster. The visualized pan-genomic data such as alignment, phylogenetic trees, maps mutations within that cluster to the branches of the tree, infers loss and gain of genes on the core-genome phylogeny for each gene cluster were presented. Besides, 997 known (MIBiG database) and 553 unknown (antiSMASH-predicted clusters and Pfam database) genes of biosynthesis gene clusters types and orthologous groups were mined in all strains. This workflow could also be used for other species pan-genome analysis and visualization. The display of visual data can completely duplicated as well as done in this paper. All result data and relevant tools and files can be downloaded from our website with no need to register. The pan-genome and biosynthetic gene clusters analysis and visualization can be fully reusable immediately in different computing platforms (Linux, Windows, Mac and deployed in the cloud), achieved cross platform deployment flexibility, rapid development integrated software package.

**Availability and implementation:** BGDMdocker is available at http://42.96.173.25/bapgd/ and the source code under GPL license is available at https://github.com/cgwyx/debian_prokka_panx_antismash_biodocker.

**Supplementary information:** Supplementary data are available at biorxiv online.

## 1 INTRODUCTION

Bioinformatics academic free softwares generally have the shortcomings of installing and configuration difficulty, large dependencies, limit size of the data uploading to online serversand so on. Therefore a lot of excellent softwares could not be fully used by biologists. Sharing bioinformatics tools with Docker can reproducibly and conveniently build all kinds of bioinformatics workflows. It gives programmers, development teams and biologists of bioinformatics the common toolbox they need to take advantage of the use, and building, shipping and running any app, anywhere your distributed applications.

Thus, Dockers technology is very suitable for the application in the field of bioinformatics, because of its advantages and characteristics that allow applications to run in an isolated, self-contained package that can be efficiently distributed and executed in a portable manner across a wide range of computing platforms. (Belmann et al, 2015; Hosny et al, 2016; Aranguren and Wilkinson, 2015), At present, there are many bioinformatics tools based on docker are developed and published, such as perl and bioperl (Martini, 2016), python and biopython (Moghedrin *et al.*, 2016), R and Bioconductor (Eddelbuettel *et al.*, 2016), contribute their Official Docker image; famous Galaxy also contribute docker galaxy (Björn, 2016). It’s reasonable to predict that docker will become more and more extensive in the field of bioinformatics. We used docker technology to rapidly construct a pan-genome analysis process, which can be set up in the Linux, windows and MAC systems (64-bit), can also be deployed in the cloud such as Amazon EC2 or other cloud providers. The workflow will greatly facilitate biologists to apply in bioinformatics fields. Docker containers have only a minor impact on the performance of common genomic pipelines (Tommaso *et al.*, 2015)

*Bacillus amyloliquefaciens* has been researched and explored extensively and intensively with the abilities to inhibit fungi and bacteria (Nam *et al.*, 2016). [Based on the Docker we quickly run an container (on ubuntu 16.04 and win10 host) of analysis and reveal the pan-genome and biosynthetic gene clusters basic features of 44 *B.amyloliquefaciens* strains, and achieve visualizations result. The analytical workflow consists of three toolkits: Prokka v1.11(Seemann, 2014) prokaryotic genome annotation, panX (Ding *et al.* 2016) anlanysis and visualization of pan genome and antiSMASH3.0 (Weber *et al.*, 2015) biosynthesis gene clusters, which we wrote Dockerfile of BGDMdocker with all of the application and its dependencies, (we also wrote standlone Dockerfile of Prokka, panX and antiMASH in order to meet different application requirements of user. recommended in this way, see Supplement)以上也看不懂. Here we described in details how to build workflow and manipulate the analysis.

## 2 METHODS AND IMPLEMENTATION DETAILS

### 2.1 Installing docker

Docker-engine was installed on Ubuntu Xenial 16.04 (LTS) and win10 Enterprise operating system. Docker requires a 64-bit installation regardless of your Ubuntu version. Additionally, the kernel must be 3.10 at minimum. More operating systems installation see here.

Installation latest docker (docker-engine 1.12.5-0~ubuntu-xenial) on Ubuntu Xenial 16.04 (LTS) (Install Docker on Ubuntu, 2016):

To copy the following commands meant quickly & easily installing via latest docker-engine (ubuntu, debian, raspbian, fedora, centos, redhat, suse and oracle linux *et al.*, all applicable):

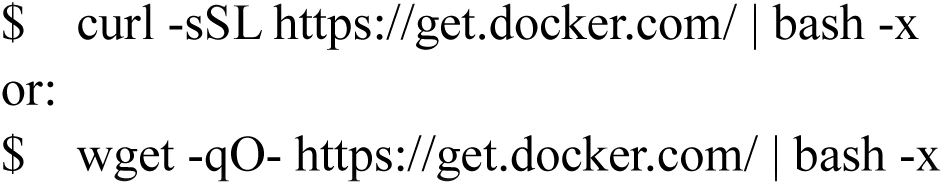

Type the following commands at your shell prompt, If output for docker version it means your installation is successful:

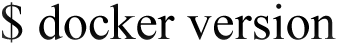

Installation latest docker on Windows 10 Enterprise(Install Docker on Windows,2016):

The current version of Docker for Windows runs on 64 bit Windows 10 Pro, Enterprise and Education.

Step 1. Download Docker InstallDocker.msi (https://download.docker.com/win/stable/InstallDocker.msi)

Step 2. Install Docker(1.12.0-rc2) for Windows

Double-click InstallDocker.msi to run the installer. Follow the install wizard to accept the license, authorize the installer, and proceed with the install.

Type the following commands at your shell prompt (cmd.exe or PowerShell), will output for docker version it means your installation is successful.

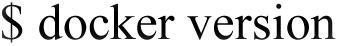

### 2.2 build images of workflow and run container login interaction patterns

Dockerfile of BGDMdocker workflow have been submitted to Github. On your host type the following commands line will login a BGDMdocker container:

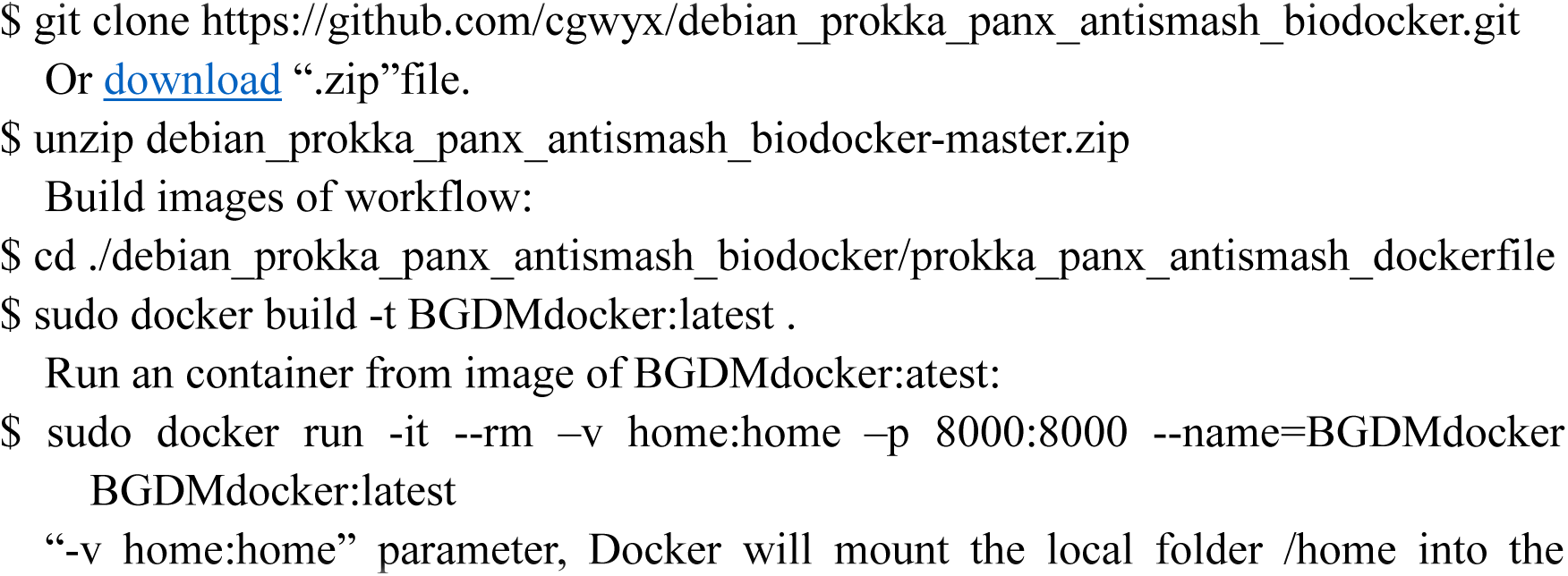

“-v home: home” parameter, Docker will mount the local folder /home into the Container under /home, Store all your data in one directory of home of host operating system, then you may access those directory of home inside of container.

Check out local images and container:

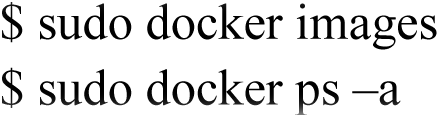

#### 2.2.1 Run Prokka annotation genome in container in interaction patterns (if you have own sequences of genome need this step generates.gbff annotation files)

Check out help documentation and command parameters:

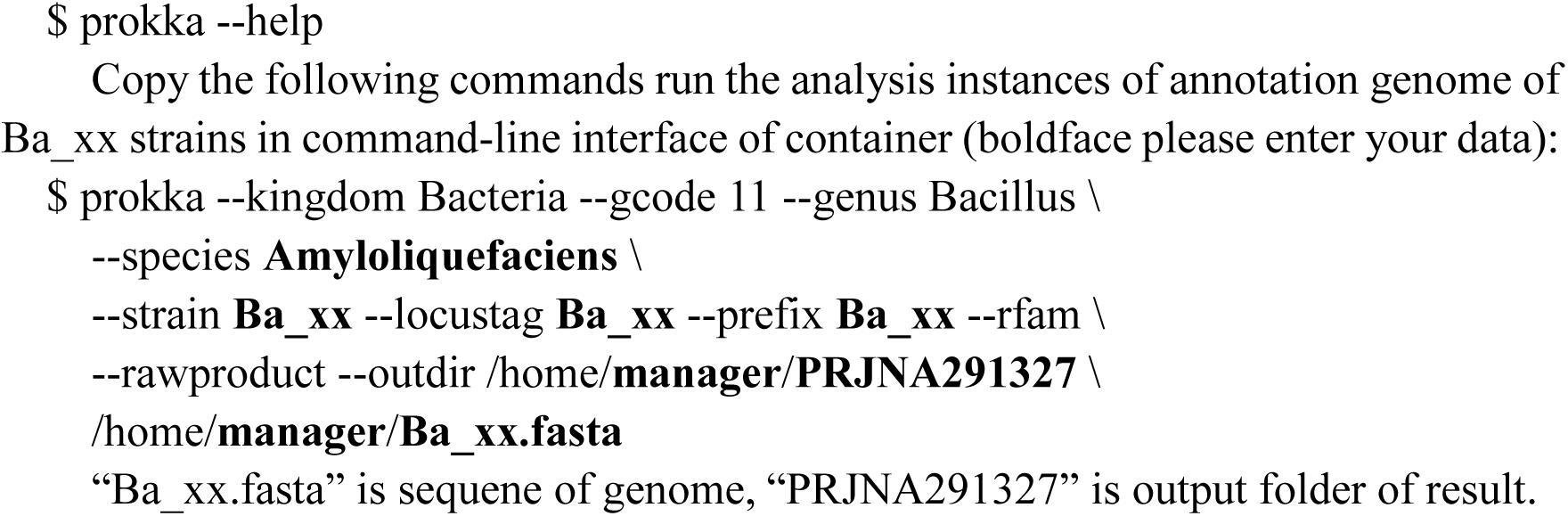

#### 2.2.2 Run panX anlanysis pan-genome in container in interaction patterns

PanX starts with a set of annotated sequences files.gbff(.gbk) (e.g. NCBI RefSeq or GenBank) of a bacterial species genomes, If using own GenBank files(or you have download these files), step 02 can be skipped. The detailed parameters see here.

Download all “*genomic.gbff.gz”of specified species from RefSeq or GenBank Database(boldface please enter your species): Installion scricp on your host

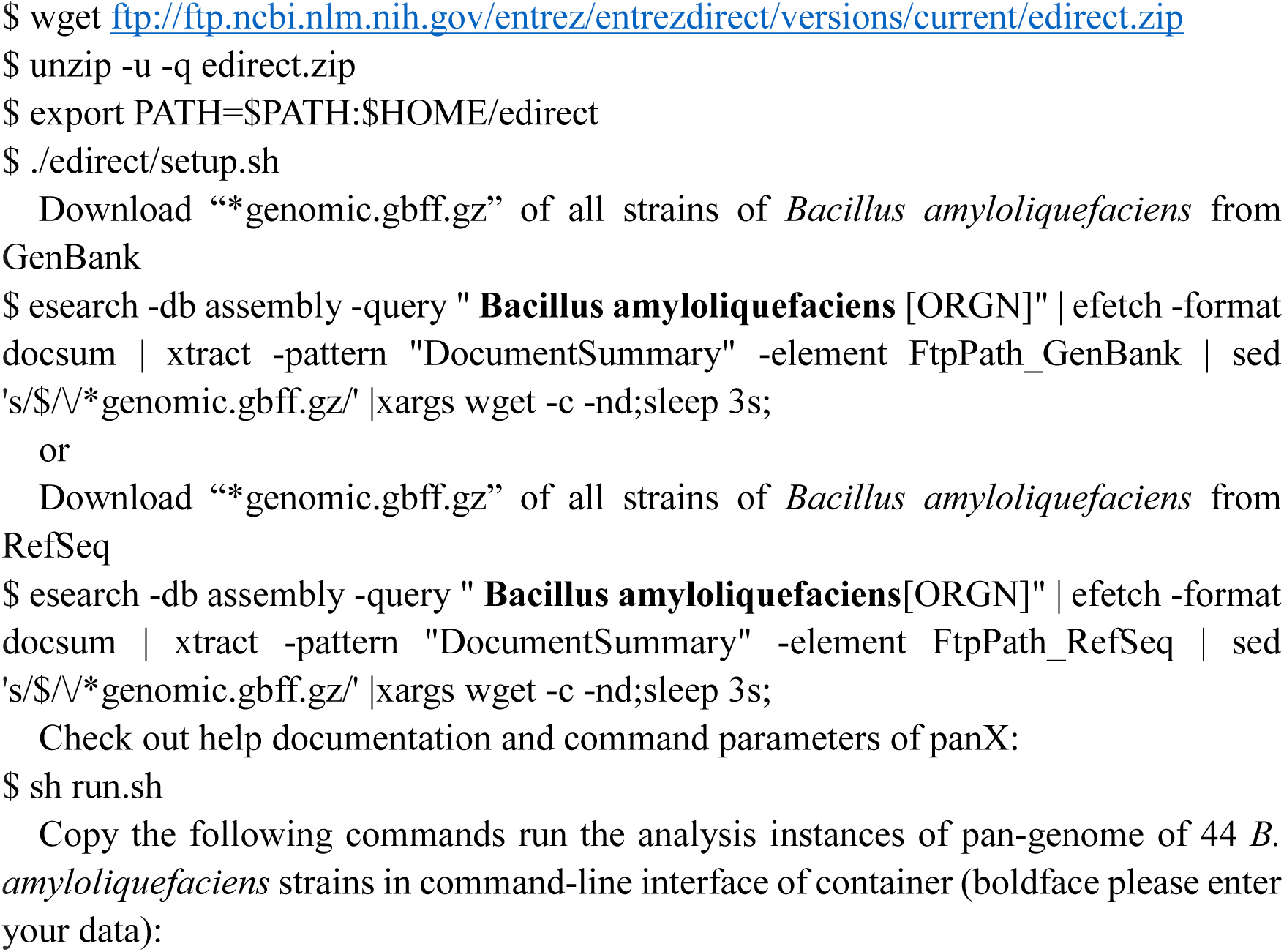

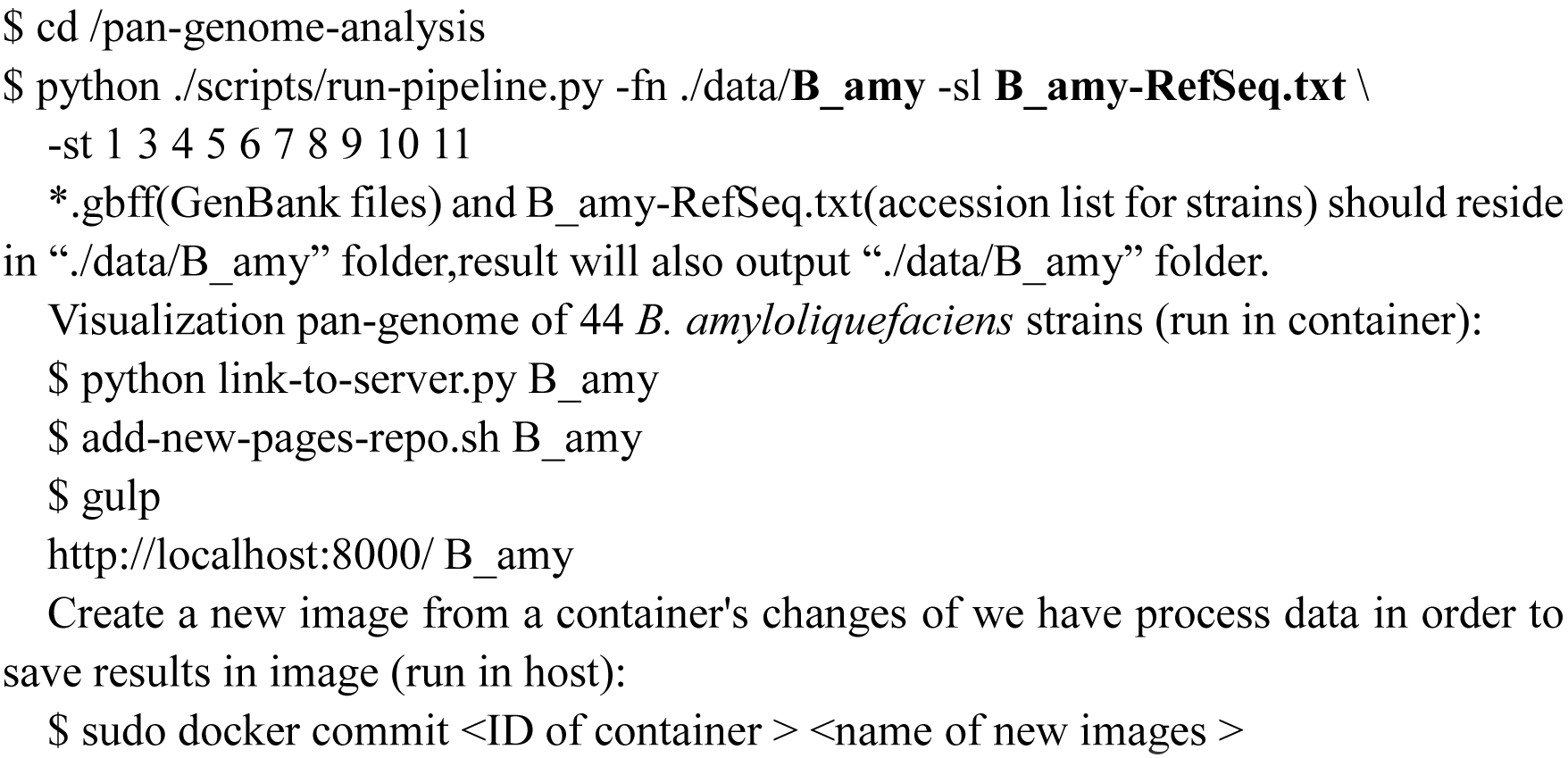

#### 2.2.3 Run antiSAMSH search gene clusters of every strain in container in interaction patterns

Check out help documentation and command parameters:

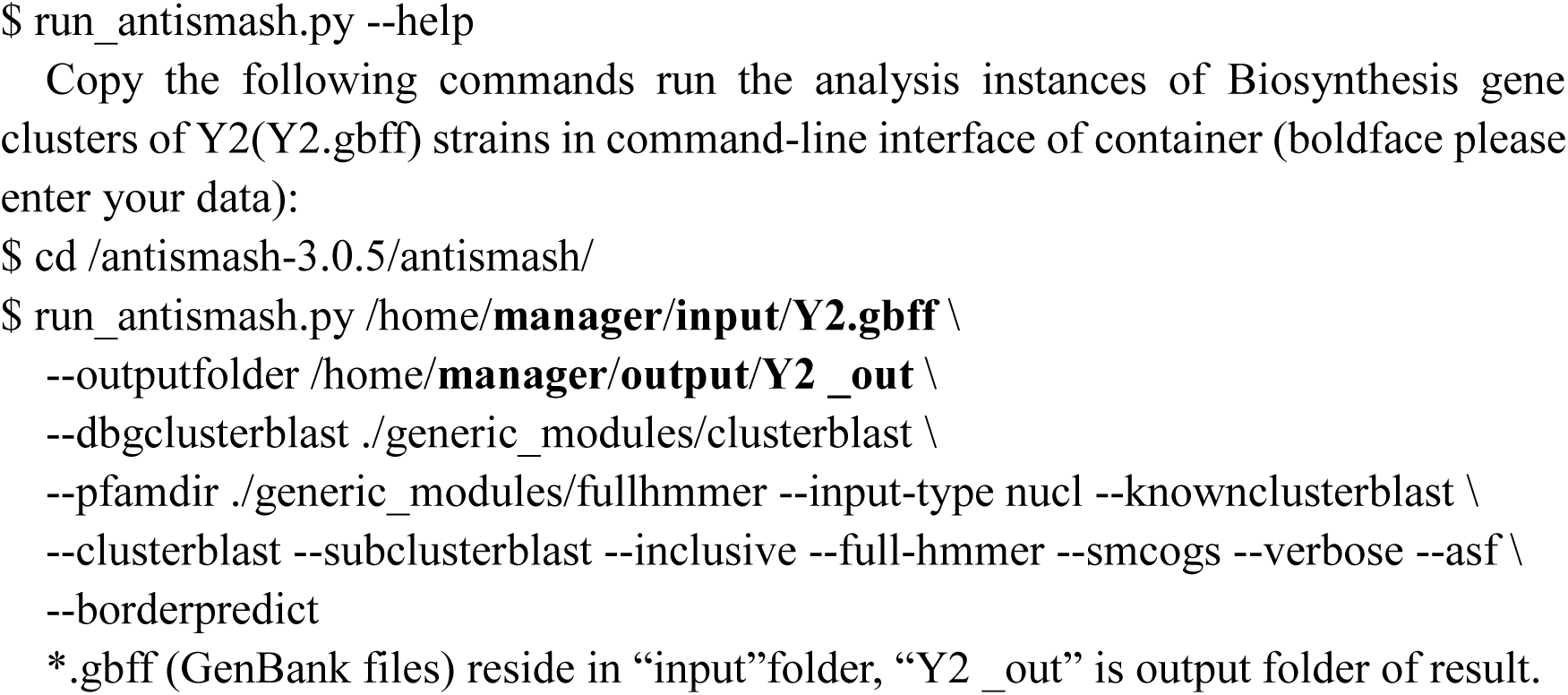

## 3 RESULTS AND CONCLUSIONS

### 3.1 Fast and repeatable buildup of BGDMdocker workflow based on Docker across computing platform

Based on docker technology, using Dockerfile script file, build images and run a containers in seconds or milliseconds on Linux and Windows (also can be deployed in Mac, cloud such as Amazon EC2 or other cloud providers). Dockerfile is just a small plain text file that can be easily stored and shared. The user does not have to deal with installing and configuring.

In this instance, all image based on Debian 8.0 (Jessie), establishment of a novel Docker-based bioinformatics platform for the study of microbes genomes. The use of workflow containers with standardised interfaces has the potential of making the work of biologists easier by creating simple to use, inter-changeable tools to excavate the biological meaning contained in data from biological experiments and Studied on the biological sequence data obtained from biological database through internet. Depending on the sequence obtained, we focused on the information mining in the sequence. We have uploaded this dockerfile to GitHub for the use of relevant scientific researchers.

### 3.2 Result of pan genomes of *B. amyloliquefaciens*

BGDMdocker workflow analyzed pan genomes of *B. amyloliquefaciens*. In order to explore high dimensional data, we built Website for interactive exploration of the pan-genome and biosynthetic gene clusters. The visualization allows rapid filtering of and searching for genes. For each gene cluster, panX displays an alignment, a phylogenetic tree, maps mutations within that cluster to the branches of the tree, and infers loss and gain of genes on the core-genome phylogeny. Here we only lists summary statistics results of pan genomes (Table 1), phylogenetic relationship of 44 *B. amyloliquefaciens* strains (Figure 1). All detail data can be visualized and downloaded without registration.

**Fig.1.**
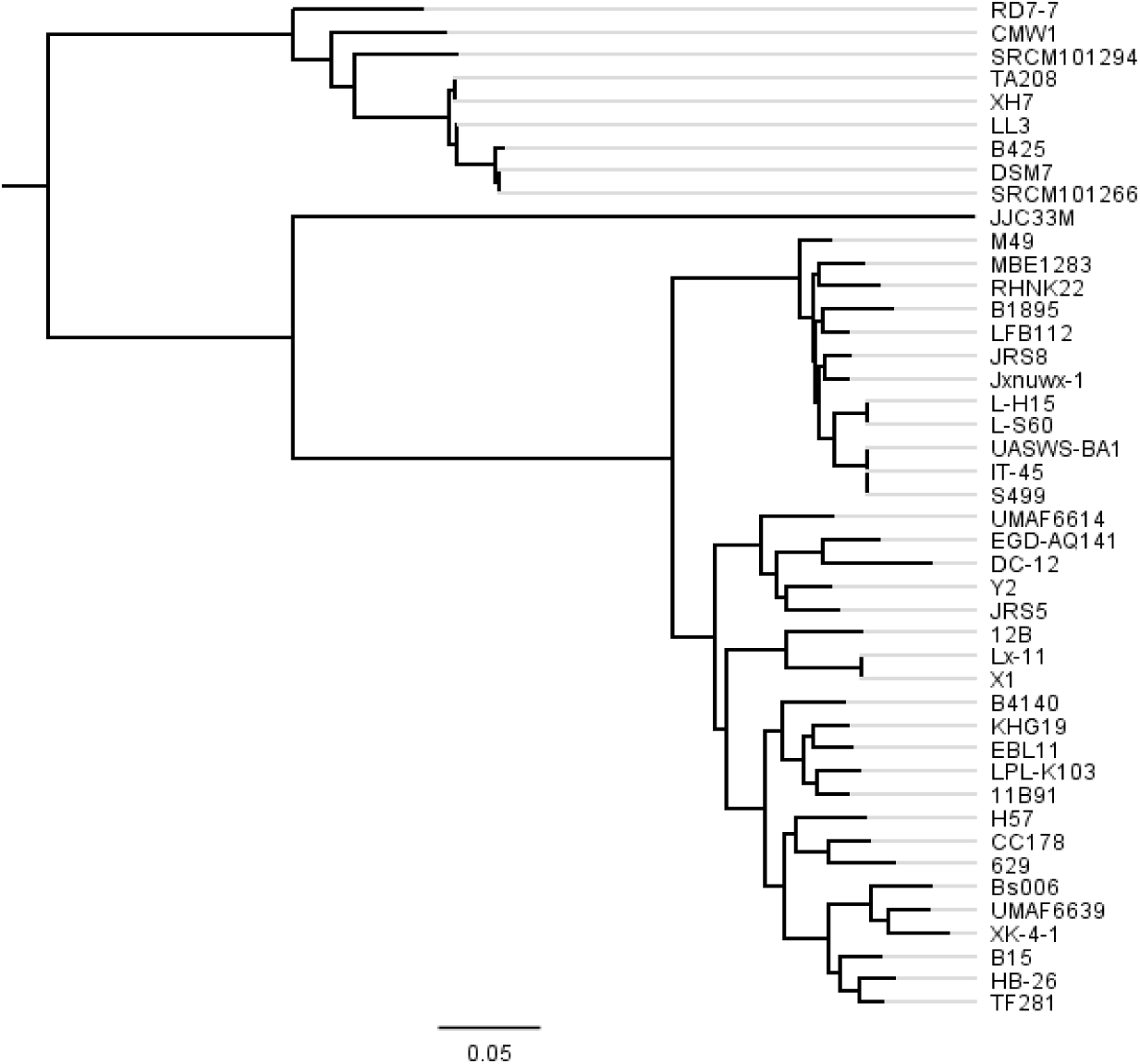
Phylogenetic tree of 44 *B. amyloliquefaciens* strains. The tree was constructed using all of the genes shared between all 44 strains (2306 core genes).

**Table 1.**
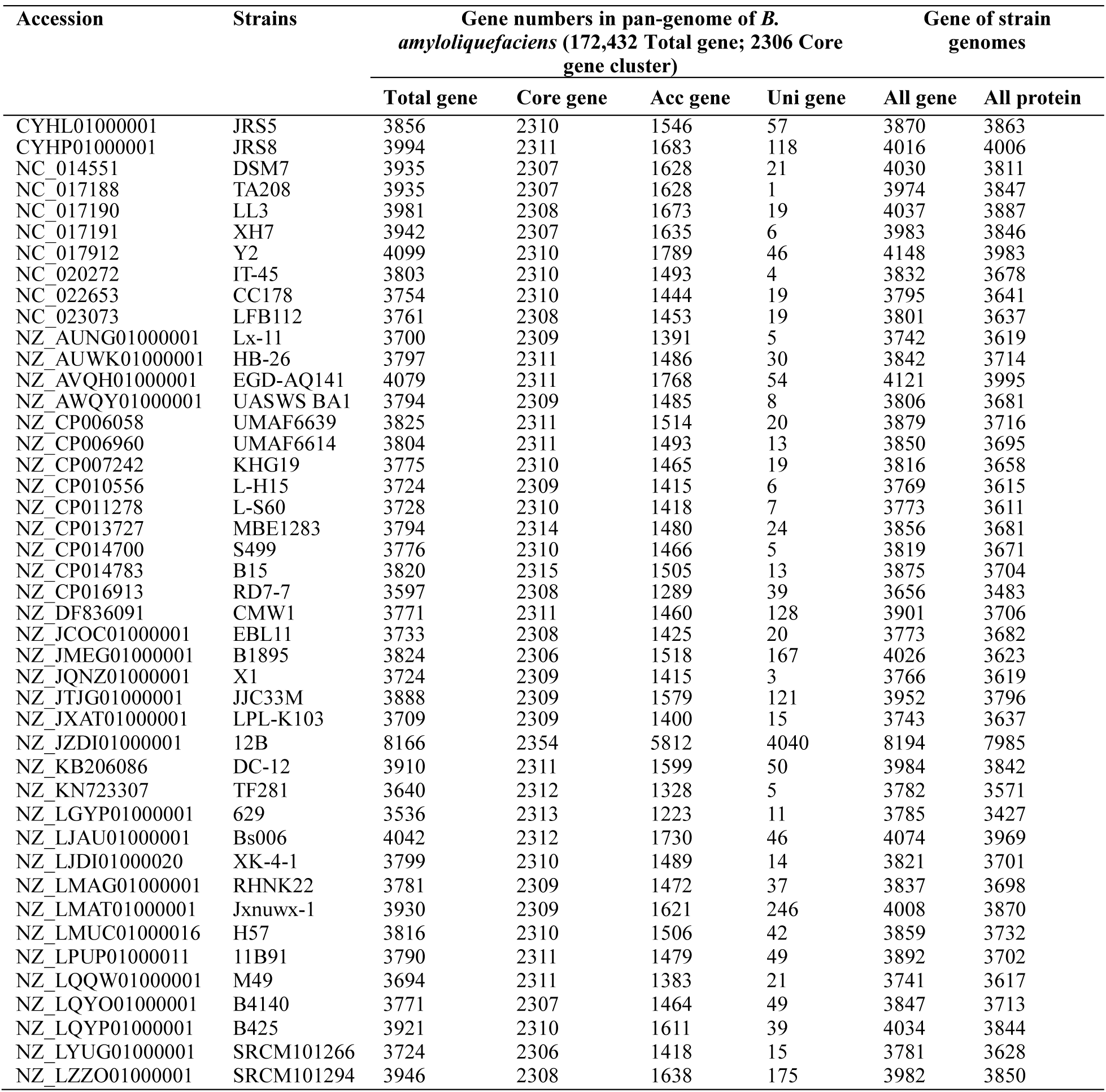
Summary statistics of pan-genome of 44 *B. amyloliquefaciens* strains

Genome sequences of 44 *B. amyloliquefaciens* strains (downloaded from GenBank RefSeq dadabase) used in this study, strains name, accession numbers, and gene numbers of pan-genome and genome summary statistics. “Acc gene”is accessory gene (dispensable gene), “Uni gene” is unique gene (strain-specific gene), “All-genes” is gene of *.gbff files recoder, incloud Pseudo Genes, “Total genes” is involved in the pan genome analysis, not Pseudo Genes.

### 3.3 Result of analysis of biosynthetic gene clusters

BGDMdocker workflow identification and analysis results of biosynthetic gene clusters of genomes of 44 *B.amyloliquefaciens* strains have been uploaded in our website, no need to register and vis and download all detail data.

In this paper we only give a brief summary statistics of biosynthetic geneclusters of all 44 strains (Table 2), and shows an instance of biosynthetic gene clusters type and number result of Y2 strain, there are total 31 gene clusters in Genome of Y2 strain. 21 gene clusters show similarity known cluster MIBiG, like this Surfactin, Mersacidin, Fengycin, and so on, other 10 gene clusters are unknown (Table 3).

**Table 2.**
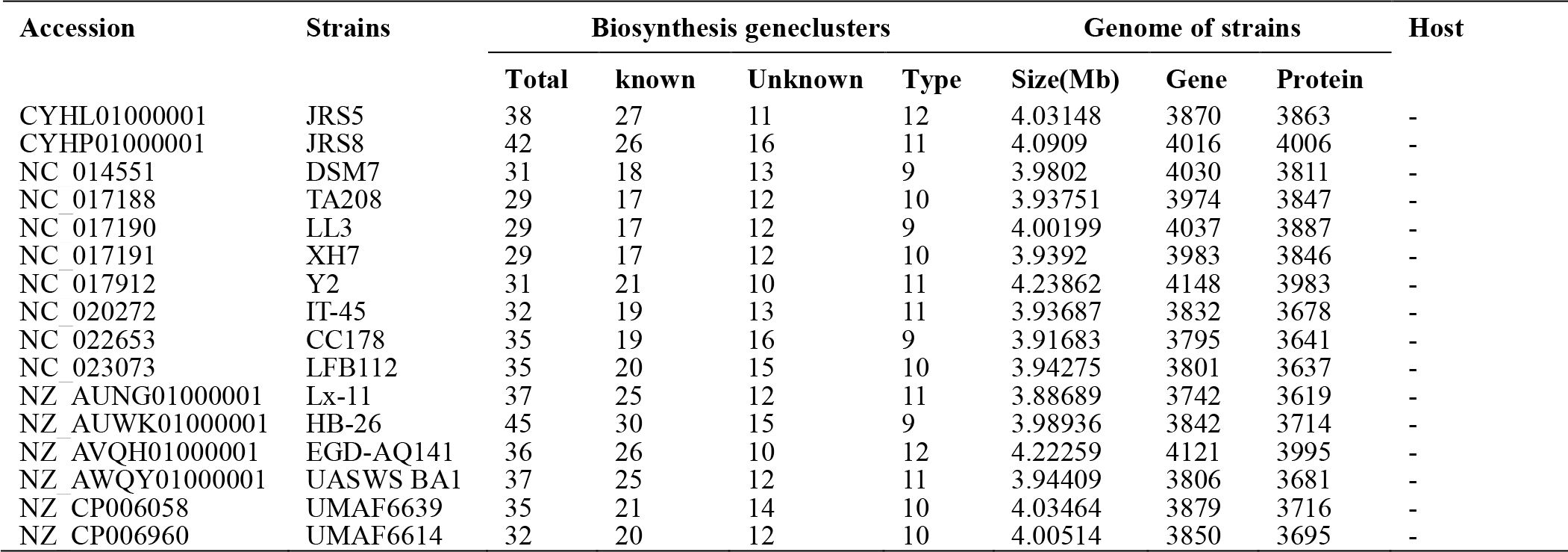
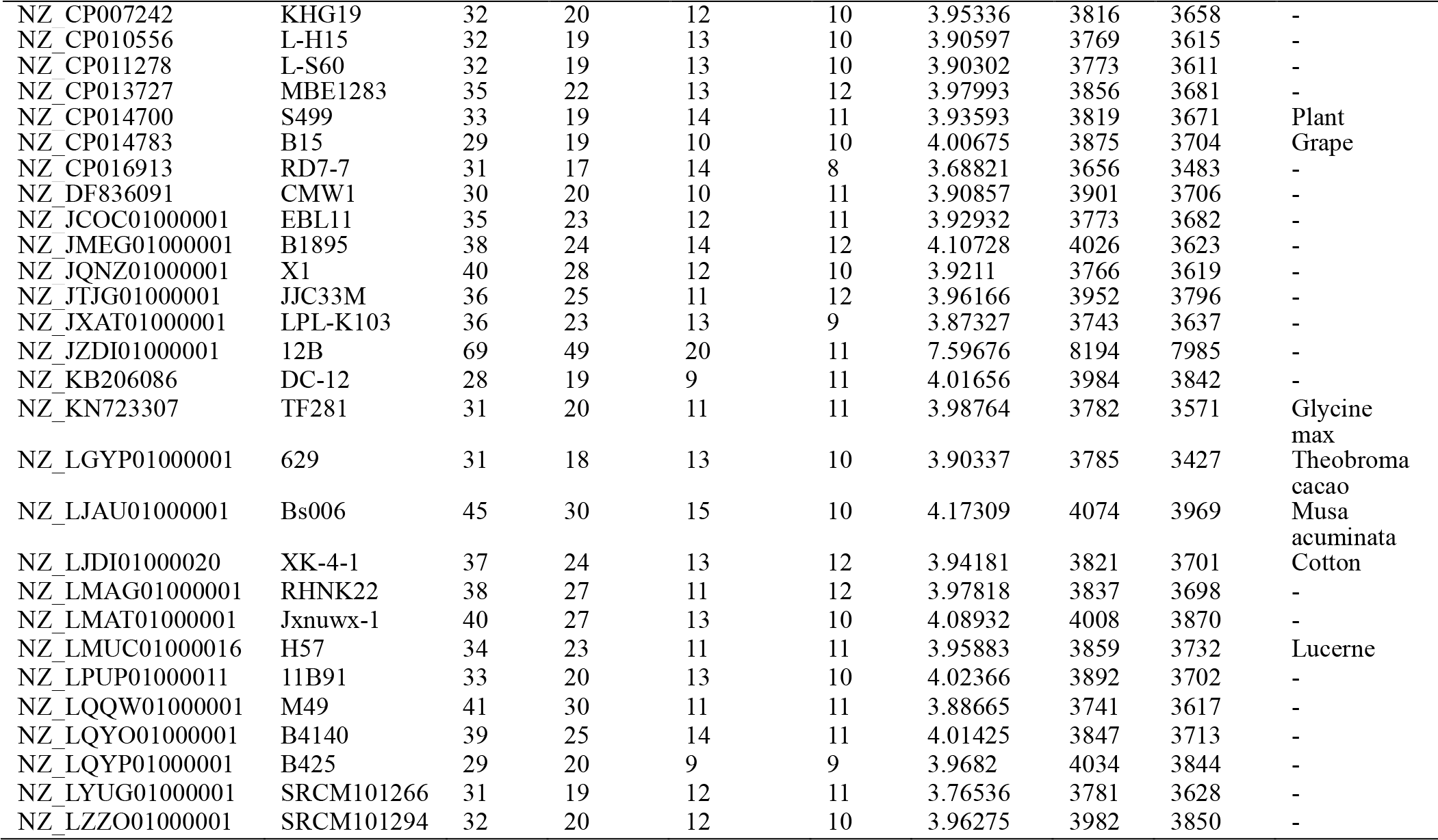
Statistics summary of biosynthetic gene clusters of 44*B. amyloliquefaciens*strains

“Total” of Biosynthesis gene clusters is include “Known”and“Unknown”. “Known” of Biosynthesis geneclusters from the MIBiG ( Minimum Information about a Biosynthetic Gene cluster).“Unknown” of Biosynthesis geneclusters detected by Cluster Finder are further categorized into putative (‘Cf_putative’) biosynthetic types. A full integration of the recently published Cluster Finder algorithm now allows using this probabilistic algorithm to detect putative gene clusters of unknown types, “-” of host is unrecorded.

**Table 3.**
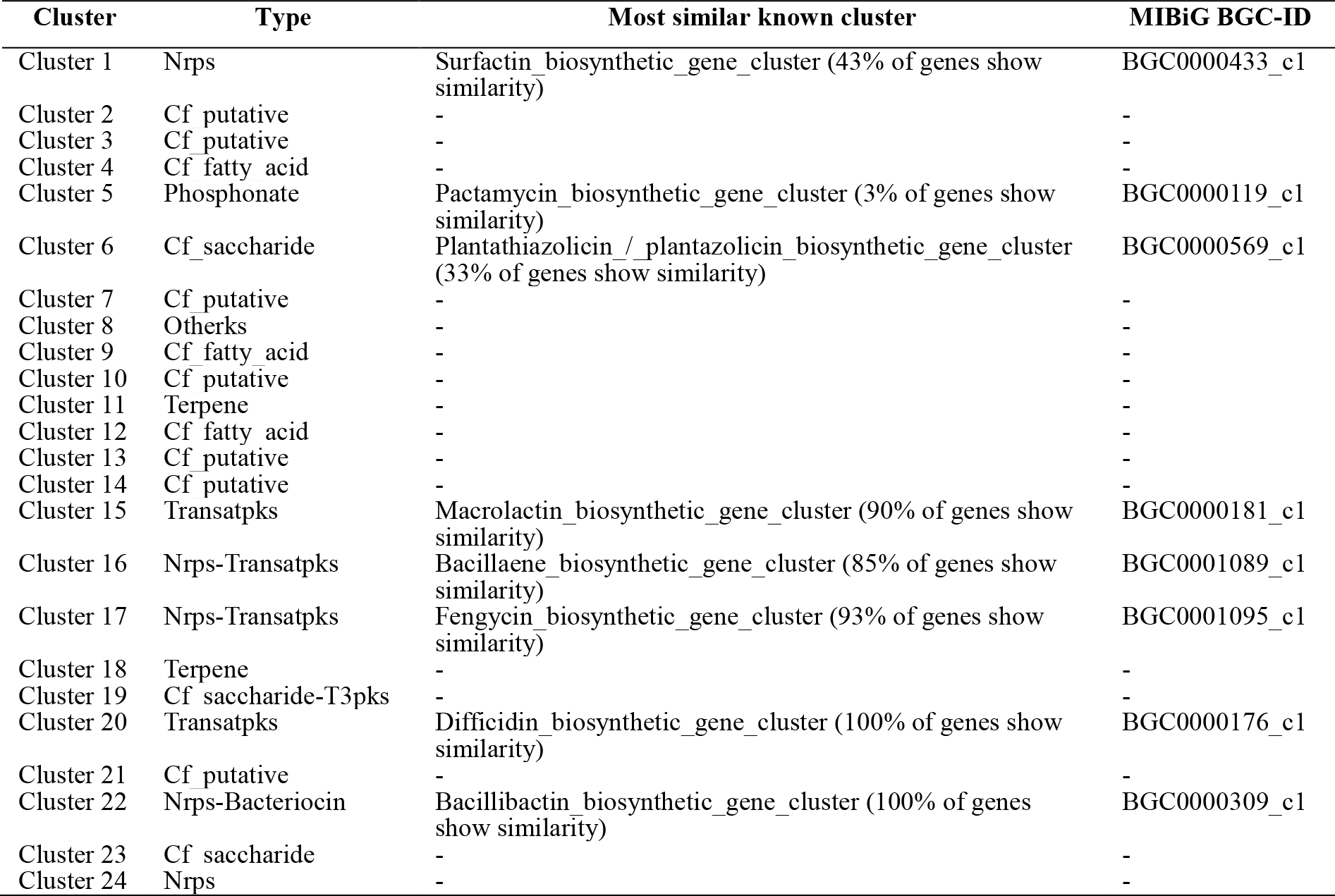
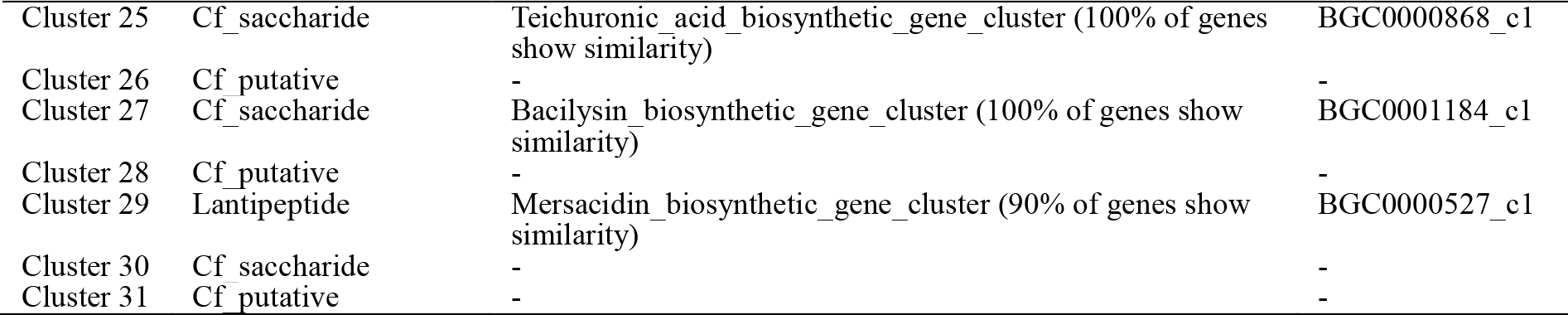
Biosynthetic gene clusters of Y2 strain

“Cf putative” is putative biosynthetic types (unknown types) detected by Cluster Finder are further categorized, known types from the MIBiG (Minimum Information about a Biosynthetic Gene cluster, http://mibig.secondarymetabolites.org).

## 4 DISCUSSIONS

Docker advantages as follow:

1. Dockerfile convenient for deployed and shared; make it is easy for other users to customize the image by editing the Dockerfile directly. It very unlikely Makefile and other installation that the resulting build will differ when being built on different machines(Boettiger, 2014). Through Dockerfile, can maintain and update related adjustment, further more rapid rollback in the event of failure of System, according to the demand to control version and build the best application environment.
2. Portability, Modular reuse; Most bioinformatics tools is written in different languages, require different operating environment configuration and cross platform, Docker provide the same functions and services in different environments without additional configuration(Folarin *et al.*, 2015), so it is possible that the results of repetition and reuse tools. By constructing pipelines with different tools, bioinformatics can automatically and effectively analyze scientific problem they are concern at of biological.
3. Application solation, efficiency, exible: Docker can be run independently containers of every applications, and Management operations (start, stop, boot, etc.) of Containers in seconds or milliseconds; we may run more than hundreds of containers on a single host (Ali *et al.*, 2016). This ensure that the failure of one task does not cause disruption of the entire process, and quickly start a new container to continue to perform this task until the completion of the whole process, improve the overall efficiency.

Docker limitations as follow:

1. Docker is limited to 64-bit host machines, then making it can not to run on 32-bit older hardware.
2. On Mac and Windows OS, Docker must run boot2docker tool in a fully virtualized environment.

